# Evaluating the value and limits of null models for Lesion Network Mapping

**DOI:** 10.64898/2026.09.23.752265

**Authors:** Martijn P. van den Heuvel, Ilan Libedinsky, Sebastian Quiroz Monnens, Jonathan Repple, Luca Cocchi

## Abstract

Lesion Network Mapping (LNM) is a method used to map symptom-related circuits in the human brain. Recent work has raised concerns about the methodological framework of LNM and the disease specificity of its derived networks, which often show high spatial overlap across unrelated conditions. Null models have been proposed to address the raised limitations^1^. As useful statistical tools in network neuroscience, they are *reference procedures*; in LNM, the normative connectome matrix *C* remains the source of the derived networks also under null modelling. Their broader role as a general solution for LNM therefore warrants evaluation. Here, we examine the proposed null models step-by-step and test them empirically across multiple datasets. We show that the proposed null-model correction effectively only replaces *C* with a transformed matrix *C′* (e.g., *C′∼C−mean(C)*), one that is directly derived from *C* and retains much of its low-dimensional structure. Consequently, null-controlled LNM maps remain shaped by the same standard set of connectome-derived patterns and continue to show repetition and non-specificity across unrelated conditions. We discuss the scope and limitations of null models for LNM, and indicate that they provide statistical reference for LNM findings, but do not establish disease specificity.

## Introduction

Lesion Network Mapping (LNM) is a functional neuroimaging method that projects lesion locations onto a normative functional connectome to identify brain circuits associated with clinical symptoms. A recent investigation of LNM raised concerns about the specificity and validity of networks derived with this method, showing substantial repetition across unrelated conditions despite being presented as disease-specific^2^. Null models have been proposed to overcome this limitation and improve the clinical applicability of LNM^1, 3, 4^. Their application has since been rapidly adopted^5, 6^.

The methodological investigation, however, noted that the observed overlap arises directly from the core steps of the method itself^2^: beneath a series of analytic steps and statistical layers, LNM is a linear projection of lesion locations *M* onto a normative connectome *C*, compactly expressed as *LNM∼∑MC*^1, 2^. Consequently, LNM outcomes are naturally constrained by the structure of *C*, with the average connectivity pattern of the matrix emerging as a dominant methodological motif across presented LNM networks, together with the other standard patterns of *C* when further contrasts are examined.

In response to the observed repetition, this overlap has been presented as reflecting a convergence of the method on “brain hubs”^1, 7, 8^. While the original study expressed reservations about this narrow interpretation^2^, changes to the statistical pipeline of LNM have been proposed to correct for the “hub-related” effect or “connectome bias”^1, 8^. Zalesky and Cash particularly proposed the use of null models^1^, with the aim of overcoming the disease-specificity limitation and improving the anatomical specificity of LNM networks.

Null models are highly valuable tools in network neuroscience^9^, making their application to LNM^1^ well-motivated and important to examine. When applied to LNM^1, 2^, these approaches suggest that several LNM circuits do not exceed corresponding null expectations. Null models remain, however, *reference procedures*: they test whether an observed effect exceeds a specified null expectation. As tools for statistical inference, they do not alter the generative source of the contrast map itself, which in LNM remains the normative connectome *C*. Updating the statistical framework of LNM may improve the statistical evaluation of LNM effects, but does not, by itself, establish that surviving effects represent condition-specific circuitry.

In this note, we expand on the discussion^2^ of null models for LNM. We examine proposed null models in context of *M* and *C* and derive that their application does not alter the central constraint of LNM: they effectively transform the used connectivity matrix *C* into an alternative matrix *C′*, one that is derived from *C* (in the simplest case approximating *Cn = C−mean(C)*), and inherit much of its low-dimensional structure, closely overlapping with the procedures of specificity testing for LNM^10^. Consequently, null-controlled LNM maps remain shaped by the limited set of distinct patterns available in *C*, rather than disentangling them from the generic structure of *C*. We apply the proposed null models across multiple empirical datasets and show that null-model correction does not fundamentally change the repetition among LNM circuits.

## Null models for LNM

LNM involves extracting functional connectivity (fc) maps for a set of lesions or brain coordinates *M* from a normative connectome dataset *C*^3, 11, 12^. The standard LNM network is derived as the sum (mean) of these individual lesion fc maps, with a tendency to converge to the matrix average of *C* when applied to heterogeneous lesion sets^1, 2^. Zalesky and Cash propose spatial null models to benchmark these LNM maps to maps generated under random conditions, building on earlier work on LNM null models with randomly sampled coordinates^3, 4^. A proposed extension^1^ introduces a constraint to the null, matching randomized lesions to regions of similar local node degree. We examined these null models in terms of *M* and *C*: the original random-lesion null model (*RL*), in which random lesion/coordinate locations are used^3, 4, 13^ and the newly proposed degree-controlled random-lesion model (*dRL*), in which random control locations are matched for local degree^1^.

### Random-lesion (RL) null

The standard *RL* null-model procedure^1, 3^ consists of (step 1) generating the LNM map of interest (*LNM∼∑MC*), (2) generating sets of an equal number of randomly sampled lesions/coordinates from all locations of the brain, (3) recomputing random LNM maps for each of these permutations, and (4) using the collection of derived random LNM maps to derive an empirical region-wise null-controlled LNM z-map and corresponding p-values, which optionally can be followed by FWE or FDR correction.

In the *RL* null model^3^ the observed LNM map is derived from the set of *m* locations of interest, given by *LNM*_*obs*_ *= ∑MC*, with *M* a lesion and set size normalized lesion matrix and *C* the connectome. Next, for each permutation, a set of the same number of control coordinates is randomly sampled from the total brain mask and projected through the normative connectome *C*; similarly, *LNM*_*random*_ *= ∑M*_*random*_*C*. This procedure is repeated for 5,000 or more iterations, yielding an empirical null distribution of *LNM*_*random*_ maps against which the original observed *LNM*_*obs*_ is benchmarked to obtain voxel/region-wise z-statistics. Technically, the resulting null-controlled LNM map is given by *LNM(v)* =*[LNM*_*obs*_*(v) − μnull(v)]* and the z-map by *LNMz(v) = [LNM*_*obs*_*(v) − μnull(v)] / σnull(v)*, where *μnull(v)* and *σnull(v)* denote the mean and standard deviation of the empirically derived null distribution of *LNM*_*random*_ at each voxel/region *v*. **Figure 1A** schematizes the LNM pipeline and its augmentation with the *RL* null model.

**Figure 1.**
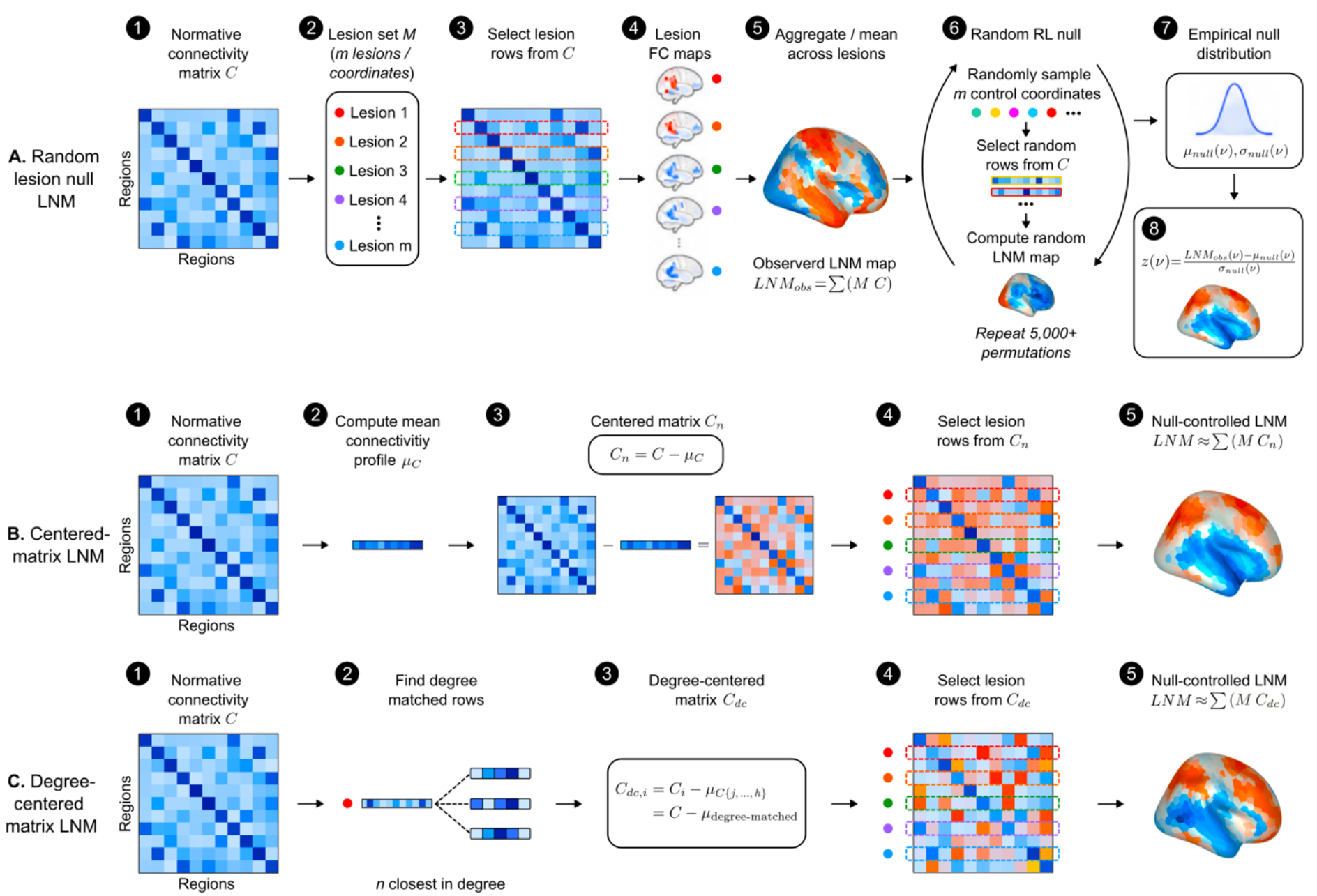
Null-model workflow and equivalent linear matrix notation. **(A)** Pipeline of the random null model (RL) variant of LNM^3^. A normative connectivity matrix C (1) is used as the reference connectome. Lesions or coordinates M (2) are projected onto this normative matrix by selecting the corresponding rows of C (3), obtaining the functional connectivity (FC) maps for each lesion (4) and aggregating these lesion FC maps into the LNM map as 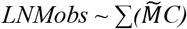, with C the connectivity matrix and 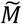 a lesion and set size normalized version of M, M for short (5). In the null-model step, for each permutation, random regions (i.e. random rows of C) are selected, and a random LNM map is estimated by means of 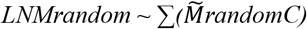 (6). Across permutations, a distribution of null effects for each voxel v is build (7), to which the observed LNM map is benchmarked, obtaining the voxelwise null-controlled LNM z-map (8). **(B)** Virtually the same map can be obtained by taking matrix C (1), subtracting the mean connectivity profile μC (2) obtaining a mean-centered matrix Cn (3) and using matrix Cn in the same procedure as standard LNM (4). This gives virtually identical maps to null-controlled LNM maps derived by the RL null model (5). **(C)** The same alternative procedure can be made for the proposed degree null model (dRL)^1^. For each row i in C (1,2), the n closest rows j … h in degree are selected (2), their μC{j…h} is computed, and used to center row Ci (3), deriving a degree-centered matrix Cdc (4). Using matrix Cdc in the LNM procedure via MCdc yields maps almost similar to those obtained by performing the degree null-permutation (5).

Similar to LNM itself^2^, this null-model component^3, 4^ allows a direct analytical formulation. With a sufficiently large null distribution, the null mean *μnull* becomes directly recoverable from *C*. Each permutation samples random rows from the voxel- or region-wise connectivity matrix *C*, and within and across permutations, the expected null map naturally converges to the mean connectivity profile of *C, mean(C)* or *μC*. Indeed, empirical sampling of random coordinates/lesions yields the pattern of *μnull ∼ μC* (r = 0.998). It can readily be obtained that the pattern of *σnull* can also be approximated directly from *C* by *∼ σC* (r = 0.997).

This implies that the contrast tested in the null-controlled LNM map can be derived analytically as *LNM ∼ [LNMobs − μC]*, with *μC* derived from *C*. Optionally, this can be scaled by *σC* to obtain values in the range of z-scores (up to a constant factor). **Figure 1B** illustrates these steps, interactive code examples are presented at nullmodels.lnmviewer.org.

An implication of this formulation is that the *RL* null model represents a centering operation on *C*. The above procedural steps are linear operations on *C*, and their order can be exchanged: benchmarking the LNM map to the null distribution is equivalent to first centering each row *i* of *C* by *Ci − μC*, deriving a new matrix *Cn = C − μC*, and then computing the LNM map directly as *LNM ∼ ∑MCn*. If desired, values can be scaled by *σC* and a constant *m* to obtain values in the scale of z-scores. Indeed, using the implementation *MCn* gives virtually identical results to applying the full *RL* permutation null model to exemplary lesion sets (r = 0.99).

This shows that the application of the *RL* null model in LNM does not expand the space of possible LNM networks, but instead retains much of the characteristic structure and patterning of *C*. Subtracting *μC* changes one dominant connectome-derived pattern from *C*—the mean connectivity profile, interpreted as the “degree” or “hub-related” pattern—but leaves the residual structure similarly constrained by the underlying matrix, leaving repetition among null-controlled LNM networks when studied across multiple conditions.

### Random-lesion degree-centered (dRL) null

The proposed *dRL* extention^1^ matches the lesions with null lesions with similar network degree locally. This is a common and useful strategy for controlling topological bias in network studies^9,^ _14_ and worth examining in the context of LNM. Accordingly, in the *dRL* model, for each examined lesion or coordinate, the *n* regions *{j,…,h}* with the most similar degree are identified (e.g. *n* = 50, 100 or 200^1^), and random locations are sampled from this subset in each permutation. Repeating this procedure again for 5,000 or more times yields a more focused degree-constrained empirical null distribution, against which the observed LNM values are benchmarked. This produces a degree-controlled (*dc*) z-map, defined as *LNMz(v) = [LNMobs(v) − μnull*_*dc*_*(v)] / σnull*_*dc*_*(v)*.

This procedure also represents a centering operation on *C*, and can be equally captured by replacing matrix *C* with an alternative matrix *Cdc* derived from *C*. For each row *i* of *C*, repeated sampling from the *n* degree-matched rows converges to the mean connectivity profile of that degree-matched set, *μC*_*{j,…,h}*_. Each row *i* can therefore be centered by subtracting this degree-matched mean, giving *Cdc*_*i*_ *= C*_*i*_ *− μC*_*{j,…,h}*_. Applying this operation row-wise yields a nodedegree-corrected matrix *Cdc*. The *dRL*-derived LNM map can then be computed as *LNM ∼ ∑MCdc*, optionally scaled by *σC*_*{j,…,h}*_ or *σC* to approximate z-score values (up to a constant derived from *m*). Indeed, testing across lesion sets (see also below for empirical results), a *MCdc* implementation produces network maps that are highly similar to those obtained from applying the full *dRL* null-model procedure (r = 0.98). **Figure 1C** further schematizes this derivation.

These formulations illustrate that null models such as *RL* and *dRL* operate by transforming *C*, effectively replacing it with an alternative matrix *C′*. But because *C′* is itself derived from *C*, the resulting null-controlled LNM z-maps remain constrained by the connectome-derived structures of *C*. The low-rank connectivity subspaces of the transformed matrices *Cn* and *Cdc* substantially overlap with those of the original matrix *C* (PC1-3 ∼ 81%), leaving the null-corrected LNM maps largely shaped by the same low-dimensional patterns of *C*. Consequently, when applied to empirical lesion sets, *RL*- and *dRL*-corrected LNM maps continue to show high repetition among unrelated symptoms and disorders and remain limited in specificity (see below).

We note that it is equally possible to express the null models through updating *M* rather than *C*: the examined contrast in the random-coordinate *RL* null model^1, 3, 4^, for example, can also be written as a direct adjustment of *M* for the number of possible regions *R* in *C, LNM ∼ (M−1/R·1)C*, which is equivalent to *M(C − μC)*. Such a notation can naturally be extended to other null models, such as the *dRL* model proposed by Zalesky and Cash^1^, where the reference expectation can be made location-specific by using a degree-matched sampling matrix *Q*, giving *M(I − Q)C*. These alternative formulations make the central point even more explicit: whether the null-model correction is expressed through updating *M* or *C*, the resulting LNM and null-controlled maps are derived through *C* and therefore remain shaped by the standard connectivity patterns contained in *C*.

### Empirical null-controlled LNM maps

We computed *RL* and *dRL* null-controlled LNM maps for 64 empirical lesion sets^15^ (Methods). Across datasets, null-controlled LNM z-maps continue to show repetition across unrelated clinical conditions (*RL* z-maps: mean |r| = 0.41, s.d. = 0.25; *dRL* z-maps: mean |r| = 0.43, s.d. = 0.25; examples shown in **Figure 2**). This level of similarity is comparable to the overlap reported for standard LNM, specificity and sLNM networks (mean |r| = 0.40^2^), with null-controlled LNM networks similarly showing |r| > 0.7 with multiple other unrelated conditions.

**Figure 2.**
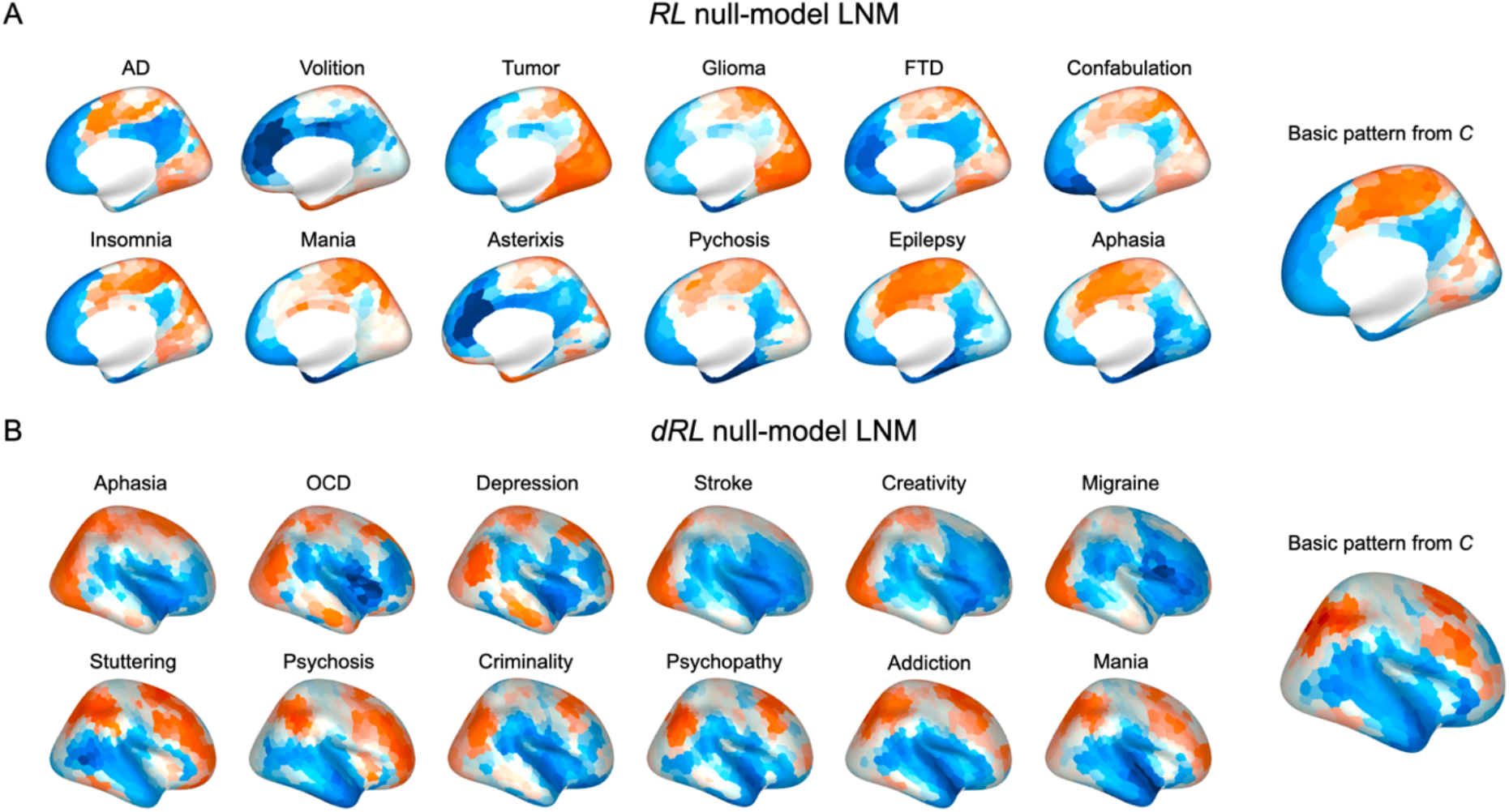
Null-controlled LNM maps retain repetitive connectome-derived structure. LNM z-maps derived by means of applying the random lesion (RL) and degree-controlled random lesion (dRL) null models. Panels show maps for exemplary lesion sets, with maps scaled to z-scores for visual comparison^2^. **(A)** Null-controlled RL maps derived by applying the RL null model to LNM, **(B)** degree null-controlled z-maps as implemented in the code of Zalesky and Cash^1^. Panels to the right show a basic pattern in matrix C, derived from fitted values of the weighted linear regression model using basic properties of C as predictors (degree, subcortical degree, modules, principal gradients (PCs) of C). We note that RL and dRL maps show overlap with standard LNM maps, suggesting that degree null correction often attenuates map amplitude rather than eliminating the underlying LNM pattern. Consequently, maps closely aligned with the dominant connectivity structure of C are mostly attenuated, whereas maps diverging from this structure are shifted toward an alternative, but still repetitive, connectivity-defined pattern in C.

We next examined the extent to which the proposed null-model corrections preserve the dominant low-dimensional spatial structure of *C* and, with it, the raised limitation of repetition among LNM maps. With *Cn* and *Cdc* transformations of *C*, subtracting *μC* from the original LNM map closely recapitulated the *RL* map across datasets (mean |r| = 0.97, s.d. = 0.008; *dRL*: mean |r| = 0.75, s.d. = 0.20). Consequently, the similarity structure among all pairs of the LNM maps remained predominantly preserved following null-model correction: conditions with similar LNM maps before correction tended to remain similar after correction (*RL*: r = 0.77 across all pairs; *dRL*: r = 0.64). This indicates that null-model correction primarily ‘shifts’ the outcome maps from one dominant connectome-derived pattern to another, rather than removing the repetitive structure.

We performed principal component analysis and fitted a simple linear model of the standard properties of *C* to the null-controlled LNM maps, further examining the lowdimensional organization imposed by *C*. Across *RL* maps, a single component explained up to 47% of the spatial variance across empirical lesion sets (first five components 93%). In addition, the standard connectivity factors derived from *C* (subcortical, cortical degree, four modules, three PC gradients^2^; **Figure 2**) accounted for on average 78% of the variance of the 64 null-controlled LNM *RL* maps. A similar organization was observed for the *dRL*-derived networks, where a single PC component explained 56% of the variance (first five components 92%), with again most of the cross-condition similarity among maps preserved (r = 0.64). As a practical example, the null-controlled LNM z-map for psychosis continued to show strong spatial similarity with LNM circuits for unrelated conditions also after *RL* or *dRL* null-control, for example with mania, insomnia, or stuttering; patterns readily found in *C* (**Figure 2**). Together, these findings suggest that disease specificity remains limited in LNM, also after null-model correction is applied.

## Discussion

Null models are central tools in neuroimaging and network neuroscience^9, 16^. Given their general utility in the field, the proposal to use them in LNM is motivated. When applied, they show that many reported LNM networks do not survive appropriate random-lesion reference procedures^1, 2^.

Earlier reservations regarding the use of null models in LNM^2^, however, were intended less as a call for the development of more constrained null procedures than as a concern about their interpretability: null models may change the comparison baseline and statistical evaluation, but they do not change the source of the contrast being evaluated. Well-designed null models are powerful tools for statistical benchmarking, but remain *reference procedures* rather than evidence, by themselves, of condition-specific circuitry. Probing lesions through *MC*—or through variants such as *MCn* or *MCdc*—yields a limited set of distinguishable spatial patterns available in *C*. When applied in LNM, they may update the statistical benchmark, rescale the maps, and/or alter which voxels or brain regions reach significance, but the central operation performed remains the projection of lesion information through the normative connectome *C*. The disease-specificity question in LNM is therefore not only whether an LNM network survives a statistical threshold, but whether it is specific when tested against clinically unrelated disorders.

This question is difficult to assess in analyses restricted to one or a small number of conditions^1^, where recurrent patterns across disorders may be missed, or surviving effects may be mistaken as disease-specific. The limited specificity of LNM becomes particularly apparent when outcomes are compared across larger collections of lesion datasets and tested against standard generic connectome-derived patterns (e.g., **Figure 2**).

The discussion around specificity in LNM points to several potential avenues for future study. Rather than assigning each clinical condition to its own ‘symptom-specific’ circuit, future work may consider the concept of *connectome pleiotropy*: testing the hypothesis that a small set of latent brain networks contribute to many cognitive functions, brain systems, and symptoms. Further integrating shared network architecture with individual data may offer a new route toward precision neuroimaging and clinically informative brain–symptom models.

## Methods

### Lesion datasets

Analyses were performed across 64 empirical lesion datasets assembled from published lesion network mapping studies and previously collected LNM resources^2^, with 10 or more cortical lesions available. Lesions were represented in Montreal Neurological Institute (MNI) space and mapped to the Schaefer1000 atlas for atlas-based analyses.

### LNM and null models

LNM maps were computed by projecting lesion information *M* through the normative connectivity matrix *C* (*LNM=*∑*MC*)^2^, and averaging connectivity profiles across lesions within each dataset. Random-lesion (RL) and degree-controlled random-lesion (dRL) null models were implemented in atlas space using the original code provided by Zalesky and Cash^1^. For each lesion set, null distributions were generated by repeatedly sampling random atlas regions, or regions matched on network degree for the dRL model, and recomputing the corresponding LNM map. Observed maps were standardized relative to the resulting region-wise null distributions.

### Matrix-based null-model implementation

Derived equivalent matrix formulations of the null procedures were evaluated (see main text for derivation). For the RL model, the mean connectivity profile of *C* was subtracted from each row, yielding the mean-centered matrix (*Cn = C −* µ*C*). For the degree-controlled model (dRL), each row *Ci* was centered relative to the mean connectivity profile of its degree-matched reference set, yielding a degree-centered matrix (*Cdc*). Equivalent null-controlled LNM maps were computed from *MCn* and *MCdc* and compared with maps obtained from the full permutation-based *RL* and *dRL* procedures.

### Cross-condition similarity and dimensionality

Spatial similarity among standard and null-controlled LNM maps was quantified using pairwise spatial correlations across the lesion datasets. To examine the dimensional structure of the resulting maps, principal component analysis was performed across LNM maps, and the proportion of variance explained by the leading components was quantified. The extent to which null-controlled maps remained aligned with dominant connectome-derived patterns was additionally assessed by relating them to canonical features of *C*, including mean connectivity, standard gradients, network modules, and principal components^2^.

## Acknowledgements

This study was funded by an ERC Consolidator grant from the European Research Council (Grant No. 101001062 CONNECT [to MPvdH]) and a NWO VICI grant from the Netherlands Organization for Scientific Research (Grant No. VI.C.241.074 BrainDiversity [to MPvdH]). JR was further supported by the LOEWE program of the Hessian Ministry of Science and Arts (Grant No. LOEWE1/16/519/03/09.001(0009)/98) and by the Deutsche Forschungsgemeinschaft (DFG, German Research Foundation, Grant No. 565437584 and 571864092).

## Data and code availability

The preprocessed normative functional connectivity time-series from the GSP1000 dataset are available from [https://doi.org/10.7910/DVN/ILXIKS] and part of the Lead-DBS toolbox. Neuroimaging data from the Human Connectome Project are available at [www.humanconnectome.org]. Example LNM maps and lesion datasets are available from GitHub [https://github.com/dutchconnectomelab/lesionnetworkmapping]. Reported lesion or LNM data are available from the referenced papers. Voxel-wise LNM was applied using the open-source Lead-DBS toolbox (https://www.lead-dbs.org/). Network analysis of the normative connectome was performed using the Brain Connectivity Toolbox. Example code of LNM and matrix implementation is presented at https://github.com/dutchconnectomelab/lesionnetworkmapping]. Code examples of null-model and specificity testing for LNM are presented at https://nullmodels.lnmviewer.org and https://methods.lnmviewer.org.

